# Towards Coordinated International Support of Core Data Resources for the Life Sciences

**DOI:** 10.1101/110825

**Authors:** W. Anderson, R. Apweiler, A. Bateman, G.A. Bauer, H. Berman, J.A. Blake, N. Blomberg, S.K. Burley, G. Cochrane, V. Di Francesco, T. Donohue, C. Durinx, A. Game, E.D. Green, T. Gojobori, P. Goodhand, A. Hamosh, H. Hermjakob, M. Kanehisa, R. Kiley, J. McEntyre, R. McKibbin, S. Miyano, B. Pauly, N. Perrimon, M.A. Ragan, G. Richards, Y-Y. Teo, M. Westerfield, E. Westhof, P.F. Lasko

## Abstract

On November 18-19, 2016, the Human Frontier Science Program Organization (HFSPO) hosted a meeting of senior managers of key data resources and leaders of several major funding organizations to discuss the challenges associated with sustaining biological and biomedical (i.e., life sciences) data resources and associated infrastructure. A strong consensus emerged from the group that core data resources for the life sciences should be supported through a coordinated international effort(s) that better ensure long-term sustainability and that appropriately align funding with scientific impact. Ideally, funding for such data resources should allow for access at no charge, as is presently the usual (and preferred) mechanism. Below, the rationale for this vision is described, and some important considerations for developing a new international funding model to support core data resources for the life sciences are presented.

## Articulating the problem

The life sciences research enterprise relies extensively upon a set of core resources that archive, curate, integrate, analyse, and enable ready access to data, information, and knowledge generated worldwide by hundreds of thousands of researchers supported by hundreds of millions of dollars of annual research investment. Some such resources are public repositories of primary data (e.g., nucleic acid sequences and protein structures), while others are public knowledgebases that assemble and curate information and insights about a particular scientific domain, organism, or biological community (e.g., communities of microbial cells). Many of these core data resources arose from modest beginnings, in some cases with histories that span more than 50 years. Some began as printed books or CD-ROMs that were regularly updated, and then morphed into web resources as the Internet became better established in the 1990s.

Today, these web-based data resources are heavily accessed around the globe by researchers in academia and industry, students and clinicians, and the interested public. They are critical for ensuring the reproducibility and the integrity of research processes [1]. The ability to deposit to and download data from these resources freely and without restrictions facilitates progress in life sciences research. Significant loss of data from these resources or introduction of barriers to data access could have devastating consequences for science, medicine, and wider society.

Core data resources are funded by a variety of mechanisms - mostly reflecting the history of how each developed over time. Some are funded by single sources and others by several sources; in almost all cases, the funding comes from national or non-profit granting agencies. The use of public funds to support this essential infrastructure ensures a strong return to society on public investments in research, and, furthermore, enables data to be reused, sometimes in unanticipated ways. However, the current funding model is fragile, with many of the data resources subject to vulnerabilities associated with grant funding, such as changing priorities, processes, and policies. Of particular concern are relatively short funding cycles (e.g., 3-5 years), and the challenges encountered when grant applications for data resource infrastructures have to compete with research proposals.

In addition, many more areas of the world are research-intensive than was the case when these data resources were first developed decades ago. In some cases, these areas are associated with substantial technical expertise that could make important contributions to operating and improving these resources. Moreover, scientists in these geographies are members of the global research community and rely on these resources in the same way as scientists elsewhere. In this regard, all life scientists, irrespective of where they are based, are stakeholders in the sustainability of core data resources.

## Defining core data resources

In order to design and implement an international plan for long-term sustainability, it is important to determine which data resources are of fundamental (i.e., core) importance to global life sciences research. This is a challenging undertaking given the scope, heterogeneity, and complexity of both the resources and the data they contain. For example, the online *Nucleic Acids Research* database catalogue lists around 1600 molecular biology data resources [2]. While some are no longer used or maintained, others have operated for decades and form a globally coordinated infrastructure that serves hundreds of thousands of researchers daily [3]. Operation of these long-standing resources requires a robust governance structure, active service management, and community-driven scientific development that are collectively well beyond the scope of a typical research program of an individual investigator. Some of these resources are connected to institutions committed to service provision [4, 5], while others have effectively navigated major management changes [e.g., transition of the Protein Data Bank (PDB) archive from Brookhaven National Laboratory to the Research Collaboratory for Structural Bioinformatics (RCSB) consortium after 27 years of operation [6]].

These long-standing data resources fall broadly into two categories:

### Archival data repositories

contain primary experimental data upon which many other databases are built. Typically, these repositories distribute data at no charge and without limitations on use, reflecting the widely held view that these fundamental data constitute a public good. Current best practices in the life sciences call for data producers to deposit primary data and metadata into such repositories prior to manuscript submission (or even sooner), with those data then made publicly accessible upon the manuscript’s publication (or before). Archival data repositories include the collection of nucleotide sequence data managed by INSCD, the International Nucleotide Sequence Database Collaboration [3], and the PDB [7], which contains information about the three-dimensional structures of biological macromolecules and is managed by the Worldwide Protein Data Bank (wwPDB) partnership. A more recently established example is the ProteomeXchange collaboration, which brings together four proteomics databases across the U.S., Europe, and Japan [8].

### Knowledgebases

add value to primary data by integrating information from multiple sources, often using computational approaches, and typically include expertly curated material. Some have a very broad scope, such as the Universal Protein Resource (UniProt [9]), which covers protein sequences and function, MetaCyc, which contains extensive information on metabolic pathways and enzymes from organisms across all domains of life [doi: 10.1093/nar/gkv1164], KBase, a collaborative open environment for systems biology modeling of plants, microbes, microbial communities, and microbiomes [doi: 10.1101/096354], and the Kyoto Encyclopedia of Genes and Genomes (KEGG), which focuses on genes and genomes [10]. More specialized knowledgebases, with deep integration of a particular domain, include the Online Mendelian Inheritance in Man (OMIM) database [11], the Arabidopsis Information Resource (TAIR), the *Escherichia coli* database (EcoCyc; doi: 10.1093/nar/gkw1003), and Model Organism Databases (MODs), such as the Mouse Genome Database (MGD), the Saccharomyces Genome Database (SGD), the Rat Genome Database (RGD), the online database of the genetics of C. elegans (WormBase), the online database for Drosophila genetics and molecular biology (FlyBase [21]), and the Zebrafish Information Network (ZFIN [12-16]). Note that the latter six knowledgebases plus the Gene Ontology Consortium (GOC [17]) recently formed the Alliance of Genome Resources (AGR; see http://www.alliancegenome.org).

The core data repositories and knowledgebases mentioned above are presented as representative examples, and are not intended as exclusionary.

## Assessing life sciences data resources

In determining whether a life sciences data resource merits ‘ core’ designation (and thus shared international support), we recommend the use of a broad set of well-defined and transparent indicators, such as those already being used by the European life science infrastructure ELIXIR [18]. These indicators are both quantitative and qualitative, with some mapping to the FAIR principles to make data Findable, Accessible, Interoperable, and Reusable [19, 20] and others measuring the impact of the resource on the scientific community and its role in accelerating science. Such indicators should also assess scientific focus and quality, the size of the research community served, the quality of the technical services provided, and the presence of a governance structure that supports open science.

While the set of data resources designated as ‘ core’ should account for long-term and international requirements, such a portfolio must be dynamic so as to adapt to changing scientific needs. In this regard, the aforementioned indicators should be used in an ongoing fashion in managing the life cycle of all core data resources – from start-up through maturity and, when appropriate, to termination.

## Determining costs and quantifying benefits

Having defined the appropriate set of core data resources for the life sciences, it will then become necessary to determine the fully burdened cost of operating each resource. In the case of archival data repositories, the replacement value of the primary data and metadata must be assessed, so as to establish whether long-term data storage is appropriate (*versus* future data regeneration on an as-needed basis). Furthermore, a reliable set of metrics for tracking the impact and cost/benefit balance of each core data resource, whether archival or knowledgebase, must be established. Finally, it will be essential to understand consequences of terminating the operation of a given resource. Addressing this final issue will require reliable quantitative and qualitative measures of the scientific, educational, and economic impact of each core data resource.

## Towards a global solution for supporting core data resources

We propose the creation of an international coalition whose mission is to collectively support those core data resources deemed essential to the work of life science researchers, educators, and innovators worldwide. Through this coalition, funders of the life sciences should commit to the long-term shared responsibility to sustain the open access to core data resources because of their value to the global life science community and adhere to the oversight principles outlined above.

The new coalition we propose would be international in scope and include representatives of major life science research funders from most, ideally all, of the countries that are active in life science research. Initial efforts of this coalition would necessarily address some guiding questions, including (1) what are the precise indicators that will be used for establishing a set of core data resources that will be eligible for shared international support; (2) will there be a binding and universal policy of global free access to the content of all designated core data resources that is appropriate and practical (as we recommend); and (3) what fraction of overall research funding from contributing nations should be dedicated to supporting core data resources (note that informal estimates of 1.5-2% have been proposed, but a more accurate accounting is warranted going forward to guide the efforts of the new coalition).

In conclusion, we believe that it is time to reshape the approach for funding core data resources in the life sciences, and we propose the launching of a coordinated international effort to harness global expertise and to create a sustainable and egalitarian data infrastructure that will support scientific endeavors well into the future.

## Disclaimer

The information and views set out in this article are those of the authors and do not necessarily reflect the official opinion of the institutions to which the authors are affiliated.

